# Deconvolution model for cytometric microbial subgroups along a freshwater hydrologic continuum

**DOI:** 10.1101/063164

**Authors:** Stefano Amalfitano, Stefano Fazi, Anna M. Romani, Butturini Andrea

**Affiliations:** Water Research Institute, National Research Council of Italy (IRSA-CNR), Rome, Italy; Institute of Aquatic Ecology, University of Girona, Spain; Department of Ecology, University of Barcelona, Spain

**Keywords:** Flow Cytometry, River Continuum, Prokaryotes, Bacteria

## Abstract

Flow cytometry is suitable to discriminate and quantify aquatic microbial cells within a spectrum of fluorescence and light scatter signals. Using fixed operational and gating settings, a mixture model, coupled to Laplacian operator and Nelder-Mead optimization algorithm, allowed deconvolving bivariate cytometric profiles into single cell subgroups. This procedure was applied to outline recurrent patterns and quantitative changes of the aquatic microbial community along a river hydrologic continuum. We found five major persistent subgroups within each of the commonly retrieved populations of cells with Low and High content of Nucleic Acids (namely, LNA and HNA cells). Moreover, we assessed changes of the cytometric community profile over-imposed by water inputs from a wastewater treatment plant. Our approach for multiparametric data deconvolution confirmed that flow cytometry could represent a prime candidate technology for assessing microbial community patterns in flowing waters.

## Introduction

Flow cytometry has been used in combination with statistical tools for dredging multiparametric representations of single cells within microbial communities from different aquatic environments (Glüge et al., 2014; Koch et al., 2014; Li, 2002). Because sample acquisition offers snapshots of single cells by delivering a multivariate dataset exportable for post-hoc analysis (Davey and Davey, 2011), different bioinformatics approaches were proposed to discriminate cytometric subgroups based on specific light scatter and fluorescence signals (Andreatta et al., 2004; Le Meur, 2013; De Roy et al., 2012). The basic cytometric cell detection combines: (i) light signals of the laser beam, scattered off at small and large angles form the cell interrogation point and related to cell size and morphology (i.e., forward and side light scatters); (ii) primary fluorescence signals related to type and content of endocellular autofluorescent pigments (i.e., autofluorescence); (iii) secondary fluorescence signals, owing to type and content of cell constitutive compounds detected upon specific staining procedures (e.g., nucleic acids) (Shapiro, 2005).

Numerous studies provided algorithms that automatically generate approximated gates to distinguish two or more cytometric groups in univariate and bivariate cytograms for a smoother interpretation of cytometric datasets (Aghaeepour et al., 2013; Hahne et al., 2009; Pyne et al., 2009; Verschoor et al., 2015). The cytometric-fingerprinting similarity among samples can be assessed and indicated through specific deviation plots and heat maps (Hsiao et al., 2016; Rogers and Holyst, 2009). However, gating and deconvolution procedures have found standardized procedures mainly for clinical diagnostic applications (Chattopadhyay and Roederer, 2012; Mittag and Tárnok, 2009; Perfetto et al., 2006).

Such procedures are fraught with failure when exploring cytometric profiles of environmental microbial communities, since the cytometric description of a natural system can be far more puzzled than that of a clinical specimen (Hyrkas et al., 2015; Koch et al., 2013b). The aquatic microbial communities comprise large populations of phylogenetically and phenotypically dissimilar cells, whose structural and cytometric dynamics depend on their specific metabolic preferences and abilities to cope with local environmental conditions (Koch et al., 2014). Moreover, dispersal of microorganisms among communities (e.g., passive movements), species sorting (e.g., selection of species within the local pool) and biotic interactions (e.g., resource competition, grazing activity, prey-predator balance) may fundamentally affect the community structure and assembly processes (Shade et al., 2012).

Natural abiotic ranges and gradients were reported to determine the cytometric fingerprinting of local communities within a given water mass at different temporal and spatial scale (Schiaffino et al., 2013; Van Wambeke et al., 2011). Moreover, a large body of literature reported a high level of analysis at log-scales to deal with the cytometric complexity of environmental samples, such as those from marine, freshwater and groundwater systems (Amalfitano et al., 2014; Boi et al., 2016; Vila-costa et al., 2012).

Given such structural and functional complexity, cytometric fingerprints may provide information on structural dynamics of microbial communities, by detecting the modifications of scatter and fluorescence signals of recurrent localized cytometric subgroups. Although disregarding a direct taxonomic sense, localization and signal intensities of cytometric subgroups have been used to define the cytometric profile of samples, which can be then compared to others from the same system. This approach can be especially effective to assess structural dynamics of microbial communities in highly dynamic systems such as flowing waters, which receive inputs from tributaries of distinct characteristics to that of the main stem. When waters flows directionally, each section along the hydrologic continuum acts as both recipient and source of waters with definite physical, chemical and biological characteristics (Nelson et al., 2009). The mass transport and the rate of external inputs to the recipient volume determine the intensity of the mass effects and the residence time available for microbial life processes (Niño-García et al., 2016).

By disregarding external inputs and stressors, it is assumable that the water network in a river system behaves as a passive corridor, particularly at high flow velocities and over short distances (Butturini et al., 2016), with the aquatic microbial community showing preserved structural traits. In such conditions, the cytometric fingerprinting might be also recurrent. Here, we provide a methodological procedure suitable to deconvolve cytometric bivariate datasets into *n* subjacent subgroups. Specifically, we tested a method for processing microbiological patterns in the headwaters of a Mediterranean river, assuming that significant external water inputs may potentially affect community structure and, thus, the cytometric profiles along the river continuum.

## Methods

### Study site and sampling

River waters were sampled during a morning survey from the upstream area of the River Tordera (Barcelona, Spain), approximately every 3 km from its natural spring to the coastline. The anthropic impact is relatively low since only small urban settlements are located within the study area. Thus, the river flows almost unimpacted for about 20 Km, until the outflow of a small Waste Water Treatment Plant (WWTP) reaches the main stem (Freixa et al., 2016). We collected five samples from the river before the WWTP outflow (T1, T2, T3, T4, T5), the WWTP waters before the conjunction with the river (A1) and the river waters after the WWTP outflow (T6). All samples were immediately fixed (2% formaldehyde, final concentration).

### Flow cytometry and cytograms

The aquatic microbial community was characterized within one week from sampling by using the Flow Cytometer A50-micro (Apogee Flow System, Hertfordshire, England) equipped with a solid-state laser set at 20 mV and tuned to an excitation wavelength of 488 nm. The volumetric absolute counting was carried out on fixed samples, stained with SYBR Green I (1:10000 dilution; Molecular Probes, Invitrogen) for 10 min in the dark at room temperature. The light scattering signals (forward and side scatters) and the green fluorescence (530/30 nm) were acquired for the single cell characterization. Thresholding was carried out using the green channel. Samples were run at low flow rates to keep the number of events below 1000 events/s (Gasol and Moran, 2015). The total number of prokaryotes was determined by their signatures in a plot of the side scatter vs the green fluorescence. The intensity of green fluorescence emitted by SYBR-positive cells allowed for the discrimination among cell groups exhibiting different nucleic acid content and morphology. The instrumental settings were kept the same for all samples in order to achieve comparable data (Prest et al., 2013).

The Apogee Histogram Software (v89.0) was used for data handling and visualization. A preliminary gating was applied to distinguish the single-celled prokaryotic community from background caused by suspended abiotic particulate, cells in aggregates and electronic noise. This step was applied to all samples during the data acquisition in a cytogram of SSC versus Green fluorescence, in accordance with previous published protocols (Gasol and Moran, 2015). The microbial community at each site was then represented in a plot of FSC vs Green fluorescence at the image resolution of 1024×1024 pixels. For each sample, the data matrix of the two variables was exported (.csv files) and deconvolved by the methodological approach described below.

### Deconvolution approach and model description

According to the Finite Distribution mixture Modelling (FDM), the complex surface *f_(x,y)_* of a bivariate cytogram (i.e., Sybr Green fluorescence vs forward scatter) is described as the sum of *n* subjacent peaks (eq. 1):

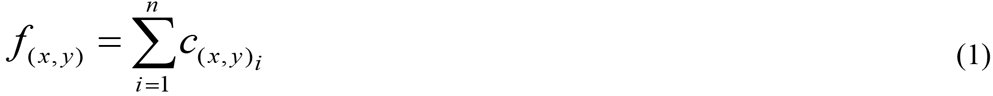

Each peak represents a subgroup that fits a predefined probabilistic density functions (*c*_*(x,y)*_). Here, an asymmetric parameter (*r*) was incorporated into the Gaussian PDF probability model (Fruhwirth-Schnatter, 2006) in order to cope with asymmetries and long tails (Kato et al., 2002) (eq. 2):

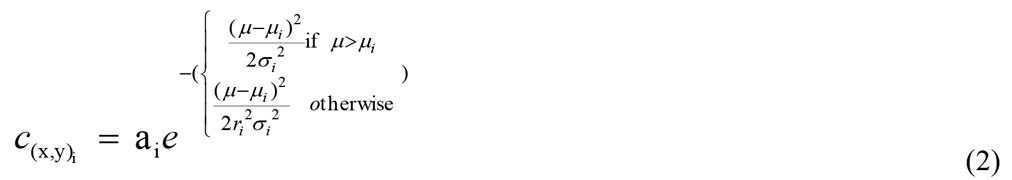

If the skewness *r_i_* (*r_ix_*, *r_iy_*) is equal to the unity, equation 2 is equivalent to a Gaussian distribution defined by its mean *μ_i_* (*μ_ix_*, *μ_iy_*), deviation *σ_i_* (*σ_ix_*, *σ_iy_*), and height *a_i_*(*a_ix_, a_iy_*).

A two-steps procedure was performed to estimate the unknown parameters of recurrent peaks (*μ_i_*, *σ_i_*, *a_i_* and *r_i_*), and to cluster those peaks into subgroups of events with a direct quantification of their density.

In step A, a surface analysis was performed according to the Nelder-Mead optimisation algorithm to detect and locate the position of local maxima (*L_n_* = {*μ_1_*, *μ_2_*, *μ_3_*, *μ_n_*}) in the *f_(xy)_*, and the position of local minima of the differential Laplacian operator of *f_(xy)_* (*∇*^2^*f*) (Ganzha and Vorozhtsov, 1996; Horst and Pardalos, 2013). To avoid overestimating the number (*n*) of potential subjacent peaks (*L_n_*), we first extracted all *i* distinct subsets of *L_n_*, (given *i*=2^*n*^−1), then we run equations 1 and 2 for each subset *i*.

In step B, the optimal number of peaks (*L_n_*) was found at the lowest value of the Bayesian Information Criterion (*BIC*) (Schwarz, 1978). To avoid meaningless results, all selected peaks must have a positive height (*a_i_* > 0). Steps A and B are detailed below.

The analysis of *f_(xy)_* was performed to detect the position of potential peaks in a density plot. This step combines two search strategies:

a. Detection of global and local maxima in the *f_(xy)_*:

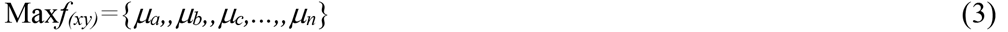
b. Detection of local minima (*μ’_i_*) of the differential Laplacian operator of *f_(xy)_* (*∇*^2^*f*):

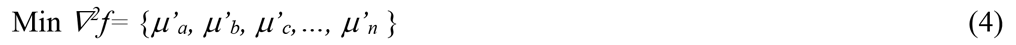

*∇*^2^*f* describes the sum of the second derivative of *f_(xy)_* with respect to *x* and *y* (Ganzha and Vorozhtsov, 1996). It was used to detect shoulders, edges and non-evident peaks in complex surfaces (Butturini and Ejarque, 2013).

The search for maxima in *f_(xy)_* and minima in *∇*^2^*f* was performed with the Nelder-Mead optimization algorithm under constrained conditions (Horst and Pardalos, 2013). The sensitivity of this algorithm can be increased or reduced by modifying selected parameters (namely: the contraction ratio, the expansion ratio, the reflection ratio and the shrink ratio). In our application, we used the standard values for these parameters (0.5, 2, 1, and 0.5 respectively) (Nelder et al., 1964) as they guaranteed an exhaustive search of main local minima in *∇*^2^*f* into a relatively short computational time. The *∇*^2^*f* operator is sensible to edges. Therefore, the minimum in *∇*^2^*f* surface found in the proximity perimeter of the cytogram were omitted. Once Max *f_(xy)_* and Min *∇*^2^*f* were obtained, results were joined to sort all distinct coordinates that appear in the two lists:

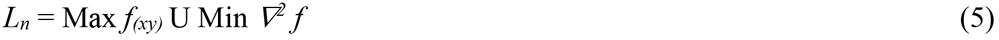

In which *L_n_* is the list of the putative *n* peaks in*f_(xy)_*.

In complex surfaces such those of cytograms, the Nelder-Mead algorithm can be trapped in local minima (or maxima), which are very close to each other and presumably identify the same peak. From a statistical perspective, it was assumed that these neighbour peaks fall into the same cluster. In this case, it is necessary to merge them into a single coordinate. The search for clusters was performed according to the *fixed radius near neighbour* approach (Bentley et al., 1977). At each detected coordinate (*μ_i_*), a circular influence area (*IA_i_*) of radius *R* is associated (*IA_i_*=*πR*^2^), centred at the point *μ_i_*. The value of the radius *R* was the same for all detected *μ_i_* and fixed to set the *IA* value to 7.5% of the planar area of the surface matrix. The coordinates within the area *IA_i_* of *μ_i_* were automatically grouped into the same cluster. Two criteria were established to assign a coordinate to each cluster:

Criterion # 1 (applicable for the equations 3 and 5): the coordinate with the highest maxima was selected, the rest was discarded.

Criterion # 2 (applicable for the equation 4): the coordinate with the lowest *∇*^2^*f* was selected, the rest was discarded.

Each cytogram was converted into an *n* × *n* array (bins of 5% of width) to obtain the surface *f_(xy)_* and its *∇*^2^*f*. When ignoring *a priori* the number cytometric subgroups, Step A identifies a set of *n* potential peaks and their coordinates (*L_n_*). In order to obtain the optimal number of subgroups, we adopted the Bayesian Information Criterion (*BIC*) descriptor:

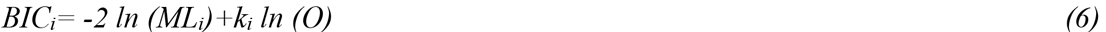

Where *ML_i_* is the maximized likelihood of the model associated to the subset *i, k_i_* the number of input parameters (i.e. number of element in the subset *i*), and *O* the sample size. The model with the smallest *BIC* value was selected as the most representative one (Schwarz, 1978). This procedure first requires the identification of all *i* distinct subsets of *L_n_*:

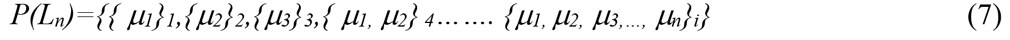

Where *i*=2^*n*^−1.

Successively, FDM (see equations 1 and 2) is run for each *i* subset, and the optimal model (i.e., the one with the optimal number of subgroups) was selected by relying on the lowest *BIC* value.

Figures 2 and 3 describes the entire process for the cytogram T6. In this cytogram the Nelder-Mead algorithm detected 3 local maxima in the *f_(xy)_* (Fig. 1a) and 7 local minima in the *∇*^2^*f* (Fig. 1b). According to eq. 5, *L_n_* represented a list of 10 potential peaks (n=10) and 2^*10*^−1 = 1023 subsets were generated (Fig. 2). The model with the lowest *BIC* values was the one with 8 peaks (*BIC*=5469, r^2^=0.978, Fig. 2). The model output with the higher r^2^ (0.979) was discarded because of the higher *BIC* value (5522). This process was executed for all cytograms in the data set.

**Figure 1.**
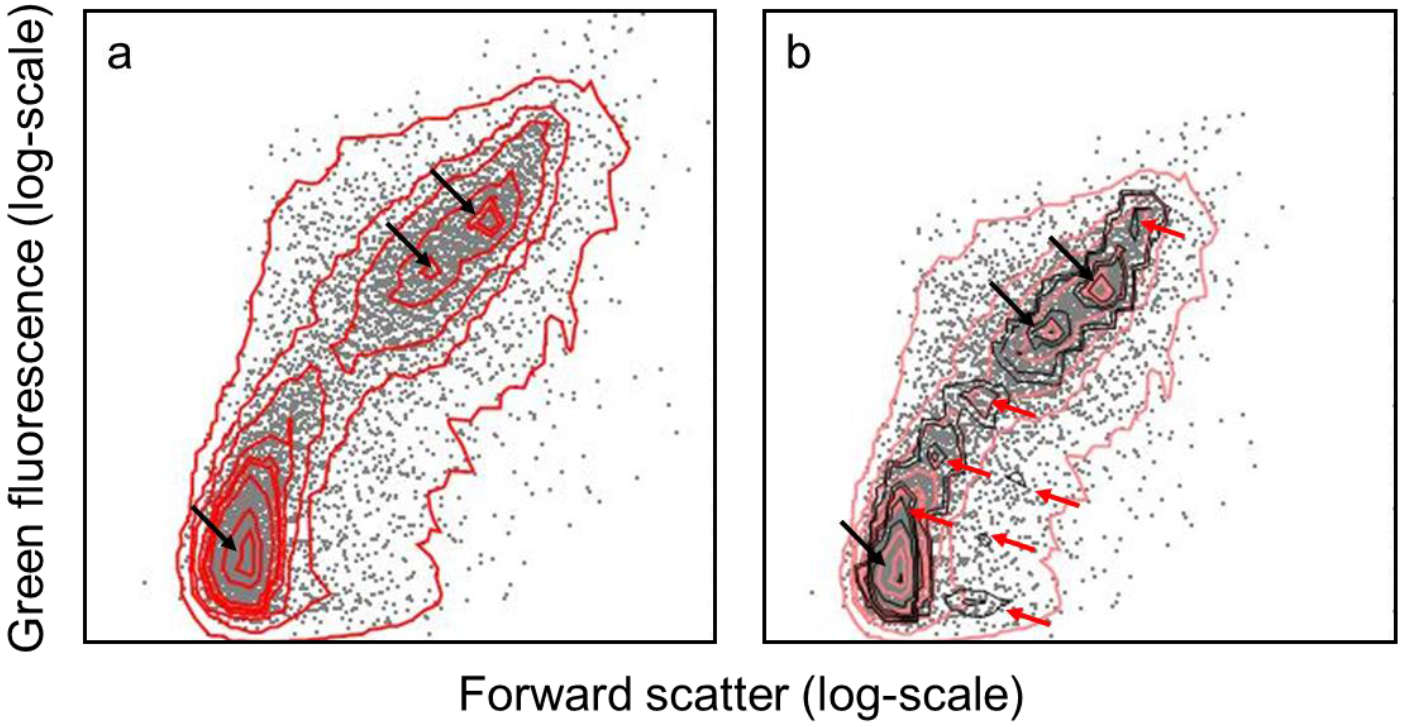
Quantile contour plot (*f_(xy)_*) of the model cytogram (T6) used to describe the deconvolution process (a) and its associate Laplacian (*∇*^2^*f*) (b). Black arrows indicate the position of the local maxima in *f_(xy)_*. Red arrows indicate the position of the local minima in *∇*^2^*f*. According to eq. 5, ten relevant peaks were detected in this cytogram from sample T6 (see fig. 3).

**Figure 2.**
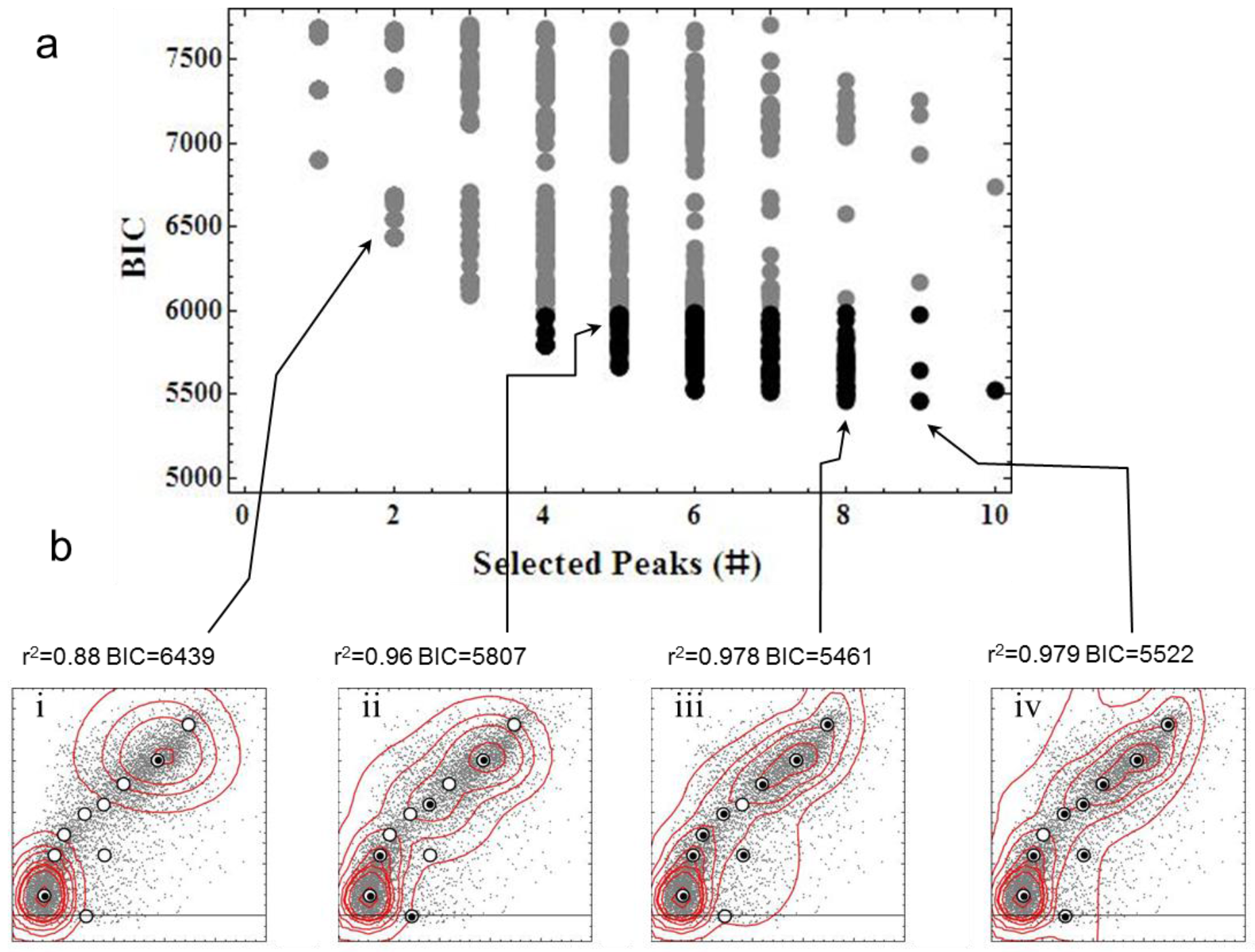
Visual example of the optimal model selection process. This example refers to the sample T6 with ten potential peaks (*n*=10) (see fig 3). Panel *a* shows the relationship between BIC values and number of peaks obtained executing the FDM *z_(x,y)_* (eqs. 1 and 2) for all possible subsets *i* of the ten potential peaks, where *i*=2^10^−1=1023. Gray disks and black dots discern modeled cytograms adjust with r^2^ lower and higher than 0.95 respectively. Panel *b* shows the contour plots of four modelled cytograms with a “poor” adjust (*i*); a good not “optimal” adjust (*ii*); the “optimal” adjust (i.e., lower BIC values, *iii*) and overfitted adjust (i.e. larger number of peaks, *iv*). Large white and small black dots in contours plots show location of potential and selected peaks respectively.

**Figure 3.**
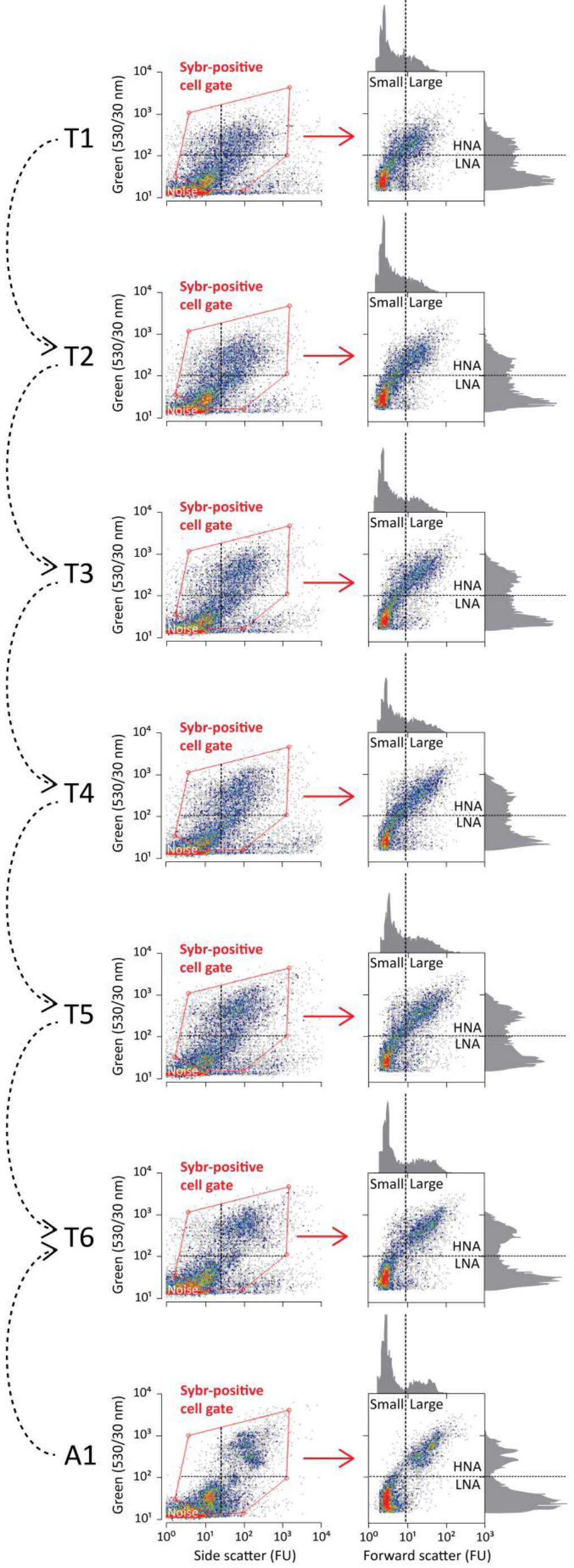
Representative cytograms of freshwaters sampled form the upstream area of the River Tordera (Barcelona, Spain). Curved arrows indicate the directional connections between sampling sites along the hydrologic continuum. The green fluorescent signals (Sybr Green I) were used to discriminate two major populations of cells with low and high content of nucleic acids (LNA and HNA, respectively). Small and large sized cells were distinguished according to forward scatter signals.

A hyper-scatterplot was created by including all *BIC* selected peaks (*μ_i_*) which were retrieved from the bivariate cytograms in the dataset.

The Voronoi diagram tessellation approach was adopted to cluster all *BIC* selected peaks into adjacent polygons with boundaries outlined by the Delaunay triangulation algorithm (Aurenhammer and Klein, 2000). All events that lie within a polygon are assigned to the centre of that polygon, and the number of events lying within each polygon was converted into cell concentration values.

### Statistical analyses

A hierarchical clustering produces, based on Ward’s method and Euclidean distance, was used to show how sampling points were clustered according to percentages of cytometric groups over the total events in the cytogram. The overall significance of such difference was tested by the non-parametric Kolmogorov-Smirnov test to give information if the densities of different cytometric groups change differently along the river system (difference in mean and if this depend on site). The multi-group SIMilarity PERcentage test (SIMPER), using the Bray-Curtis similarity measure (multiplied by 100), was run to assess which cytometric clusters were primarily responsible for the observe difference between groups of sample and the average dissimilarity among sites and (Clarke, 1993).

## Results and discussion

In line with the consolidated approach for analyzing planktonic prokaryotes across a wide range of natural and engineered water systems (Bouvier et al., 2007), two major cytometric populations, namely cells with low nucleic acid content (LNA cells) and cells with high nucleic acid content (HNA cells), could be discriminated from our river continuum samples without further data processing and counted by using fixed polygons. As expected, HNA cells were relatively brighter in fluorescence and bigger in size than LNA cells (Fig. 3). In a previous study, drinking waters were distinguishable from one another based on the percentage of HNA cells and the direct comparison of their green fluorescence histograms (Prest et al., 2013).

Our methodological approach allowed deconvolving five major recurrent subgroups within either LNA (clusters 1-4 and 10) or HNA (clusters 5-9) cytometric populations. Samples form T2 and T4 retained the lowest and larger number of subgroups, respectively (2 LNA+ 4 HNA and 4 LNA + 5 HNA). The identified peaks within the Voronoi tessellation poligons, applied for cell counting, are shown in fig. 4.

**Figure 4.**
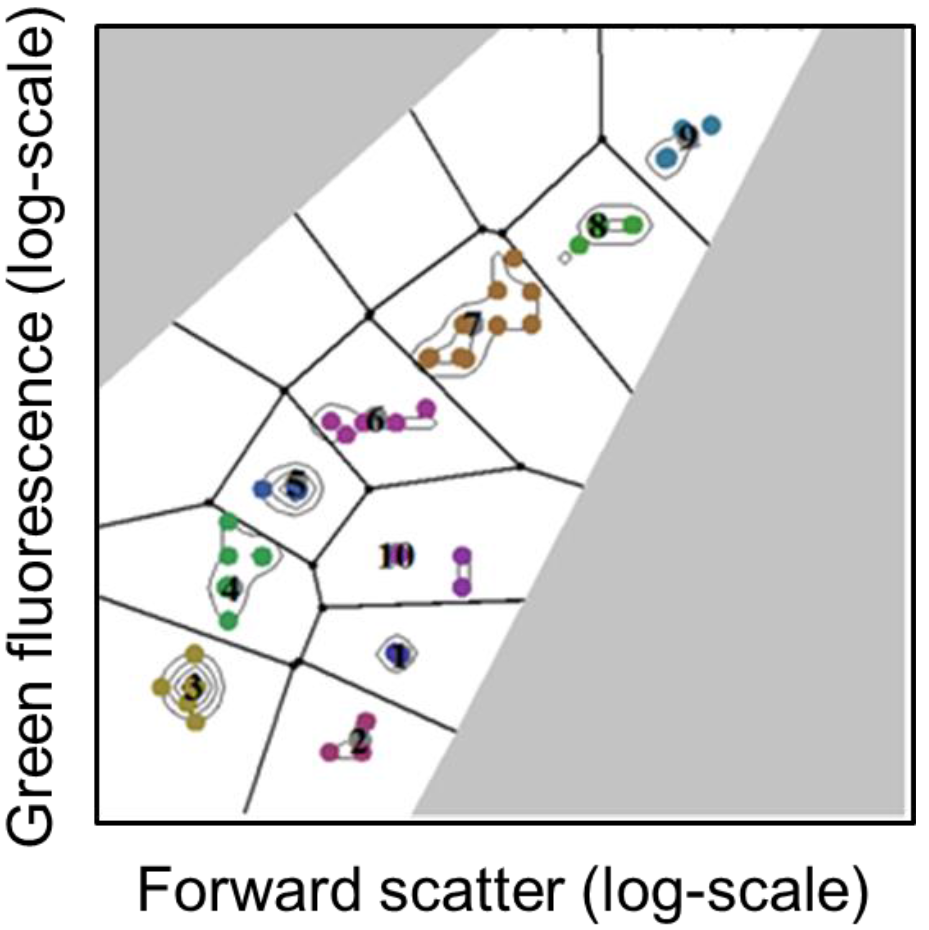
The Voronoi tessellation mask was calculated considering all recurrent peaks and applied back to each cytogram. The number of events lying within each polygon was converted into cell concentration values.

The dynamic of each identified cytometric subgroup and the evolution of the microbial community structure as a whole were assessable along the hydrologic continuum at such finer scale analysis level to provide a basis for a variety of existing statistical analysis. Hierarchical clustering offered an indication of the cytometric community similarities among sampling sites (Fig. 5). The cytometric community profiles were cross-compared (Table 1) and the community changes over-imposed by an external water input (i.e., the outlet of a wastewater treatment plant) were statistically endorsed. Moreover, clusters #3 and #8, belonging to LNA and HNA respectively, were recognized as those groups mostly contributing to the overall average dissimilarity among sites (i.e., SIMPER test). In this study, the cytometric fingerprint appeared sensitive to detect the complex microbial community dynamics of flowing waters, since expected changes due to tributary inputs were clearly detected.

**Figure 5.**
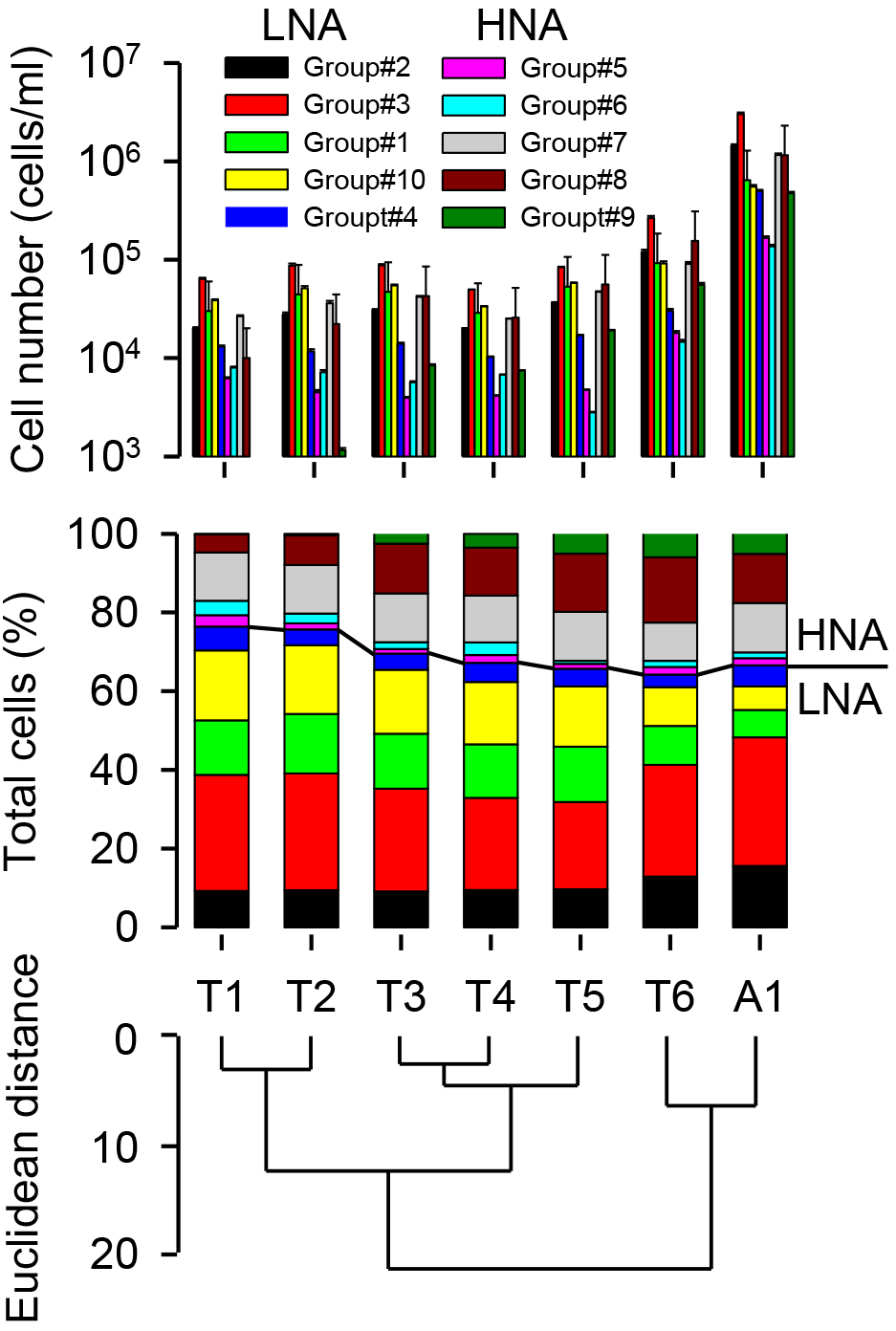
Microbial community structure as assessed by the proposed deconvolution model. a) Cell abundance within each polygon identified by the Voronoi tessellation mask. b) Relative percentages and subgroup distribution within the LNA and HNA cytometric populations. Subgroups were ordered according to their average green fluorescence. c) Hydrologically connected samples were joined according to their cytometric profile by the Ward’s clustering method such that increase in within-group variance is minimized.

**Table 1.**
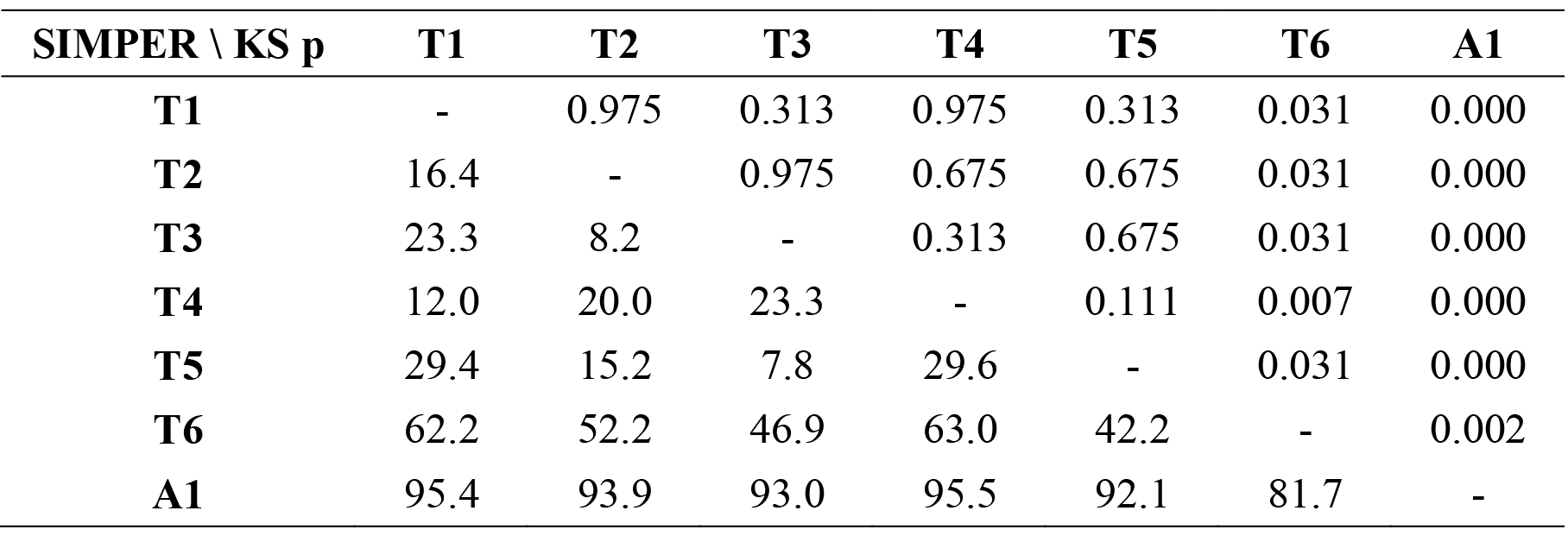
Average dissimilarity between sites computed by SIMPER test (lower part of the matrix), and related statistical diversity assessed by the non-parametric Kolmogorov-Smirnov test (upper part of the matrix).

Deconvolution models represent a way to deal with the complexity of water samples from natural and engineered water systems, allowing for an understanding of microbial interactions and structuring dynamics not described by the traditional approaches (Koch et al., 2013a).

A key perspective of the proposed deconvolution approach is the ability to discern recurrent subgroups of cells within complex mixtures, without an a priori knowledge of which cells are which. It is noteworthy that a further advanced step of cell sorting can be potentially performed to provide gate-specific phylogenetic information according to selected fluorescence properties and phenotypes of major cytometric populations (Schattenhofer et al., 2011; Vila-Costa et al., 2012).

Thousands of particles and microbial cells within a wide size range (from virus-like particles to prokaryotes and small protists) can be analyzed per second by flow cytometry, thus providing a direct quantification of their abundance and morphological traits within minutes from sampling (Van Nevel et al., 2013). A prototype machine for the automatic and programmable staining of aquatic bacteria was successfully tested on-line and in real time to monitor the quality of drinking water at the household tap (Besmer et al., 2014).

Owing to high versatility and potential for rapid analysis of large numbers of cells individually, specific benefits of flow cytometry will also accrue from novel bioinformatics and statistical approaches to analyze the multiparametric dataset, thus leading to significant and promising technological advancements in innovating the field of real time control of water quality.

## Conclusions

The application of flow cytometry to freshwater samples collected along a river continuum allowed discriminating diverse and recurrent subgroups of aquatic microorganisms by their constitutive traits at the single-cell level. Our data suggest that a flow cytometric approach could be suitable to detect changes of single-cell subgroups, thus serving as a candidate tool for water quality assessments in complex environmental settings.

## Acknowledgments

This research was partly funded by the Spanish Ministry of Education and Science (MEC) through the project FLUMED-HOTSPOTS (CGL2011-30151-C02).

